# A subset of CD4+ effector memory T cells limit immunity to pulmonary viral infection and prevent tissue pathology via activation of latent TGFβ

**DOI:** 10.1101/2023.03.02.527395

**Authors:** CP McEntee, S Houston, CM Finlay, S Rossi, G Liu, TN Shaw, J Casulli, M Fife, C Smedley, TS Griffith, M Pepper, T Hussell, PM Hansbro, J-M Schwartz, H Paidassi, MA Travis

**Affiliations:** Lydia Becker Institute for Immunology and Inflammation; Wellcome Trust Centre for Cell-Matrix Research; Division of Immunology, Immunity to Infection and Respiratory Medicine, Faculty of Biology, Medicine and Health, Manchester Academic Health Sciences Centre, University of Manchester, UK; Centre for Inflammation, Centenary Institute and University of Technology Sydney, Faculty of Science, School of Life Sciences, Sydney, New South Wales, Australia; University of Minnesota, Department of Urology, Center for Immunology, Minneapolis, MN, USA; Department of Immunology, University of Washington School of Medicine, Seattle, WA, USA; School of Biological Sciences, Faculty of Biology, Medicine and Health, University of Manchester, UK; CIRI, Centre International de Recherche en Infectiologie, ‘Normal and Pathogenic B cells’ Team, Univ Lyon, Inserm U1111, Université Claude Bernard Lyon 1, CNRS UMR5308, ENS de Lyon, F-69007 Lyon France

## Abstract

A rapid immune response to pathogen re-exposure underpins immunological memory, with protection against divergent pathogens such as heterologous or novel viral strains requiring cross­reactive memory T cells. Understanding the pathways that control memory T cell function is therefore important for the rational design of viral vaccines and will aid the discovery of therapies to boost anti-viral immunity. Here, we identify a sub-population of memory T cells that limit secondary immune responses to viral re-infection, which is crucial in preventing host tissue damage. We show that a population of CD4^+^ effector memory T (T_EM_) cells activate the important immunoregulatory cytokine TGFβ, via expression of an integrin, αvβ8. Integrin αvβ8 expression marks a transcriptionally distinct sub-population of CD4^+^ T_EM_, enriched for anti-inflammatory pathways. Loss of integrin αvβ8 on murine CD4^+^ T_EM_, but not Foxp3^+^ regulatory T cells (T_REG_), led to exacerbated virus-specific CD8^+^ T cell responses following secondary influenza A virus (IAV) infection, which was associated with enhanced viral clearance. However, although accelerating clearance, loss of integrin αvβ8 expression on CD4^+^ T_EM_ resulted in enhanced lung pathology following secondary IAV infection, which was completely reversed by adoptive transfer of αvβ8^+^ CD4^+^ T_EM_ cells. These data highlight a new pathway by which a distinct CD4^+^ memory T cell subset restrains anti-viral immunity to prevent host tissue damage during secondary viral infection. Such pathways could be targeted therapeutically to either boost memory T-cell-mediated immunity or restrain host tissue damage during viral infection.

## Main text

Whether induced by natural infection or vaccination, the development of robust immunological memory is a cornerstone of sustained protective immunity against viruses, either by directly targeting infected cells for killing or through the induction of high affinity, virus-specific antibody responses. While viral surface proteins, the primary targets for antibodies, can accumulate mutations over time leading to reduced antibody binding affinity and neutralising efficacy, memory T cells typically recognise conserved epitopes which are often shared between different viral strains or variants. Thus, virus-specific memory T cells exhibit cross-reactivity to heterologous viruses, referred to as heterosubtypic immunity, and are therefore a key component of long-lasting immunological memory, particularly as viral variants emerge over time^1, 2^. While the differentiation and maintenance of memory T cells is highly dynamic, and functional/phenotypic plasticity between subsets has been reported^3, 4^, traditionally they have been subdivided into three major subsets based on their expression of homing markers and anatomical location. Circulating CD44^hlgh^ memory T cells can be stratified into central or effector memory T cells (T_cm_ or T_Em_ respectively) based on the expression or absence of markers including CD62L (L-selectin) and CCR7 (C-C Motif Chemokine Receptor 7)^5^. CD62L^+^ CCR7^+^ T_Cm_ migrate between the circulation and secondary lymphoid structures whereas CD62L^+^ CCR7^−^ T_EM_ traffic between lymphoid and peripheral tissues. In contrast, the third major memory T cell subset reside predominantly in non-lymphoid peripheral tissues including the gastrointestinal tract, skin and lung and are referred to as tissue-resident memory T cells (T_RM_)^6^, although recent data do suggest that T_RM_ populations can also leave the tissue and circulate^7^. Given their importance in sustained, cross-reactive protective immunity against common pathogens, understanding how memory T cells function is of fundamental importance, not only to aide rational vaccine design, but also to identify targetable pathways for anti-viral immunotherapies.

Here we show that a sub-population of CD4^+^ T_EM_ control secondary immune responses to respiratory viral infection by restraining the host T cell response. These cells are specialised to activate the key immunoregulatory cytokine transforming growth factor-β (TGFβ) via expression of the integrin αvβ8, exhibit an immune suppressive transcriptional profile, and limit the frequency and cytotoxic potential of lAV-specific CD8^+^ effector T cells during a memory recall response. CD4^+^ T_EM_ expressing integrin αvβ8 directly suppress CD8^+^ T cells in a TGFβ-dependent manner and are vital in preventing tissue pathology during secondary infection. Additionally, we find that integrin αvβ8 is expressed by human CD4^+^ T_EM_, but no other memory or naïve human CD4^+^ or CD8^+^ T cells, highlighting the potential of targeting this pathway clinically to boost cytotoxic CD8^+^ T cell responses in the context of anti-viral or cancer immunotherapy. Together, our study uncovers a new suppressive role for a sub­population of CD4^+^ T_EM_ which express integrin αvβ8 and can activate latent TGFβ. These properties make integrin αvβ8^+^ CD4^+^ T_EM_ key regulators of anti-viral memory T cell responses and function to prevent inflammation-induced pathology following respiratory viral infection.

### CD4^+^ T_EM_ express *Itgbδ* and activate latent TGFβ

A crucial cytokine in the regulation of many different immune cells, especially T cells, is TGFβ^8^. TGFβ is initially secreted as a latent pro-form, consisting of the active moiety bound non-covalently and encased by latency associated peptide (LAP), from which the active cytokine must be liberated to function^9^. However, the role of TGFβ and its activation in controlling the function of memory T cell responses is poorly understood.

To investigate whether TGFβ activation may play a role in regulating memory T cell function, we first determined whether any memory T cell populations have the potential to activate TGFβ. To this end, we assessed the expression of integrin αvβ8 on memory T cell subsets, a molecule we and others have previously shown facilitates the activation of latent TGFβ by immune cells^10^. To this end, we sorted multiple CD4^+^ and CD8^+^ splenic T cell populations from Foxp3-ΎFP reporter mice^11^ to assess expression of the gene encoding the integrin β8 subunit *(Itgb8)* across several subsets by qRT-PCR (gating strategy shown in Figure S1a). As the integrin β8 subunit uniquely pairs with the αv subunit, this approach enabled us to determine relative expression of integrin αvβ8. Interestingly, in addition to the expected high levels of expression of *Itgb8* by CD4^+^ Foxp3^+^ T_REG_^12^, we also found for the first time that CD4^+^ T_EM_ (CD44^hl^ CD62L) also expressed high levels of the gene (Figure 1a). In contrast, *Itgb8* was not detected in naïve CD4^+^ T cells (CD44^l°^ CD62L^++^) nor CD4^+^ T_Cm_ (CD44^hl^ CD62L^+^). Similarly, all CD8^+^ Tcell subsets lacked expression of *Itgb8* (Figure 1a).

**Fig. 1.**
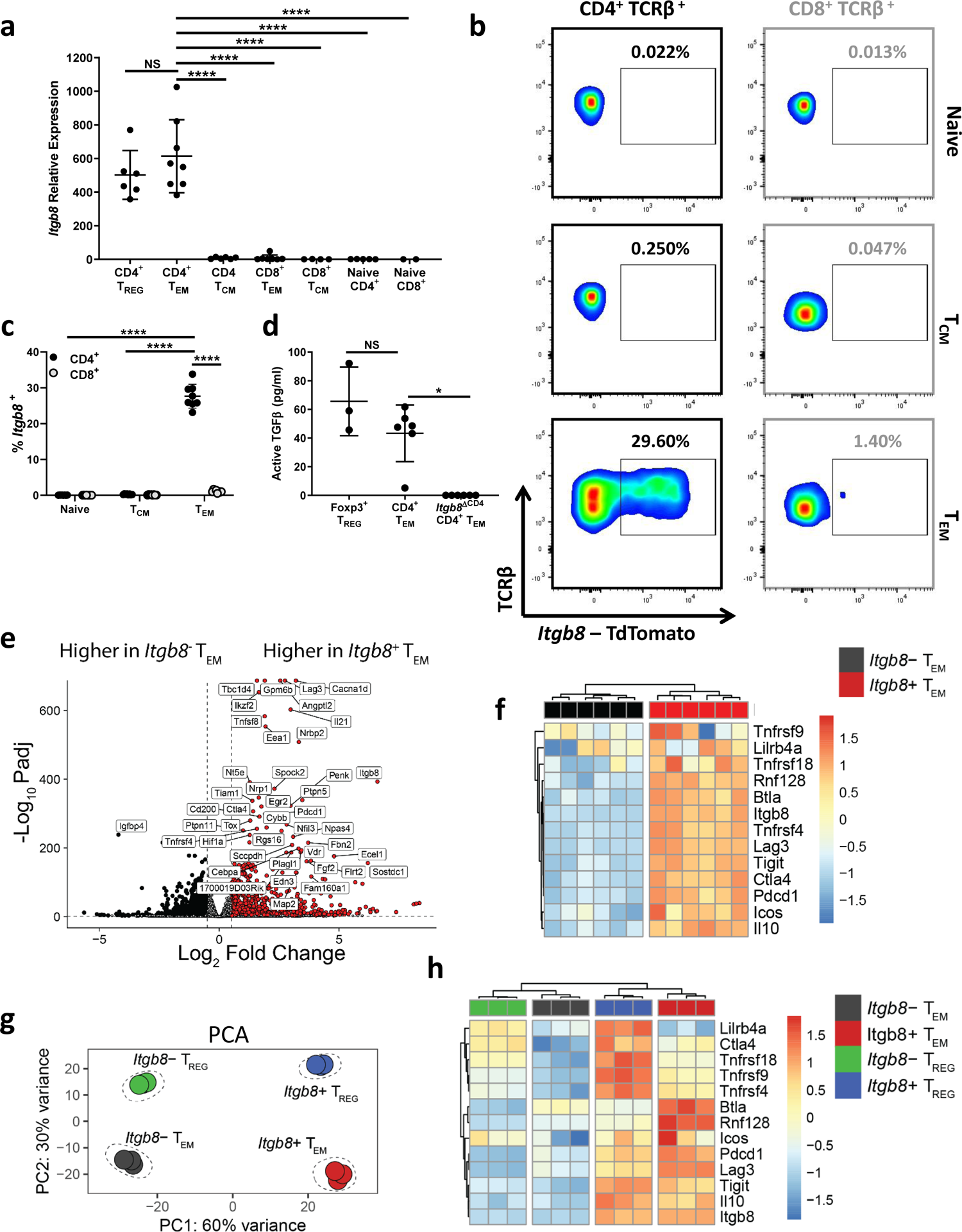
CD4^+^ T_Em_ activate latent TGFβ via integrin αvβδ and possess an immune suppressive transcriptional profile, a) Expression of *Itgb8* in FACS-purified murine splenic T cell subsets as determined by RT-PCR (n=≥6). Statistical significance determined by one-way ANOVA with Tukey’s multiple comparisons ****p=≤0.0001, ns = not significant, b) Representative flow cytometry plots of *Itgb8* expression in splenic CD4^+^ (black) and CD8^+^ (grey) T cell subsets from *Itgb8-re*porter mice, c) Collated data of *Itgb8* expression on CD4^+^ and CD8^+^ splenic T cells at steady state. Data presented as frequency of *Itgb8^+^* cells as a percentage of each parent population (n=8). Statistical significance determined by two-way ANOVA with Tukey’s multiple comparisons test ****p=≤0.0001, ns = not significant, d) Activation of latent TGFβ by T_REG_ (n=3, Foxp3-YFP^+^) or CD4^+^T_EM_ (n=6, CD44^hl^ CD62L^+^° Foxp3ļ sorted from the spleen of Foxp3^YFP^ mice or mice with a conditional deletion of *Itgb8* on T cells (/ŕgð8^ΔCD4^). Luciferase activity detected following overnight co-culture of each T cell subset with a TGFβ responsive reporter cell line. Data from two independent experiments. Statistical significance determined by Kruskall-Wallis with Dunn’s multiple comparisons test *p=≤0.05, ns = not significant, e) Volcano plot of relative expression from bulk RNA-seq (n=6) between *Itgb8^+^* T_EM_ (left) and *Itgb8^+^* T_EM_ (right), with differential (Ľog_2_ fold change > 0.5), statistically significant genes (adjusted p < 0.05) indicated in red. f) Condensed heatmap of immune suppressive genes differentially expressed between *Itgb8^+^* and *Itgb8^+^* T_EM_from bulk RNA-seq analysis (n=6). g) PCA of bulk RNA-seq experiment performed on flow cytometry-purified splenic CD44^hl^ CD62L^+^° Foxp3^+^T_REG_(/ŕgð8^+^ versus *Itgb8^+^)* and CD44^hl^ CD62L^+^° Foxp3^+^T_EM_ *Ųtgb8^+^* versus *Itgb8^+^)* (n = 3). h) Condensed heatmap of immune suppressive genes differentially expressed between CD4^+^ *Itgb8^+^* and *Itgb8^+^* T_EM_ and T_REG_ (n = 3).

Next, to assess the expression of *Itgb8* at the single cell level, we utilised a novel dual reporter mouse expressing GFP and tdTomato under control of the *Foxp3* and *Itgb8* promoters respectively^13^ (hereafter referred to as *Itgb8* reporter mice). We observed expression of *Itgb8* in a subset of CD4^+^ T_EM_, accounting for ∼30% of the total CD4^+^ T_EM_ pool (Figure 1b-c). In contrast, no expression was detected in CD4^+^ naïve or T_Cm_ cells, or any CD8^+^ T cell subsets (Figure 1b-c), consistent with our qRT-PCR data (Figure 1a). Additionally, we also observed the presence of /řgðS-expressing CD4^+^ T_EM_ at other anatomical locations, including within non-lymphoid tissues such as the lung (Figure S1b-c) and intestine (Figure S1d-e). Thus, the expression of *Itgb8* is unique to CD4^+^ T_EM_ amongst the circulating memory T cell pool.

Having determined that CD4^+^ T_EM_ express *Itgb8, we* next investigated whether this enabled these cells to activate latent TGFβ. To test this, we isolated CD4^+^ T_EM_ from mice lacking *Itgb8* expression on T cells *(Itgb8^f¦¤×,’f¦¤×^ ×* CD4-Cre, hereafter called *Itgb8^ΔCD4^*) or their littermate controls, and co-cultured them with an active TGFβ reporter cell line as previously described^12^. We observed that CD4^+^ T_EM_ activated latent TGFβ to levels comparable with CD4^+^ Foxp3^+^ T_REG_ (Figure Id). This ability to activate latent TGFβ was entirely dependent on integrin αvβ8, as TGFβ activation was completely abrogated in CD4^+^ T_EM_ cells isolated from *Itgb8^ΔCD4^* mice (Figure Id). Taken together, these data show for the first time that CD4^+^ T_EM_ uniquely express *Itgb8* amongst the circulating memory T cell pool and can activate latent TGFβ in an integrin αvβδ-dependent manner.

### *Itgb8^+^* CD4^+^ T_Em_ exhibit a transcriptional profile associated with immune suppression

Although CD4^+^ T_EM_ express *Itgb8* and can activate latent TGFβ, it is noteworthy that *Itgb8* expression was not ubiquitous but rather restricted to a sub-population of CD4^+^ T_EM_ (Figure 1b-c). These data could indicate stochastic expression within the population, or that expression of *Itgb8* by CD4^+^ T_EM_ is linked to a distinct phenotype or function compared to their ItgbS-negative *(Itgb8^+^)* counterparts. To investigate further, we purified *Itgb8^+^* and *Itgb8^+^* CD4^+^ T_EM_ from *Itgb8* reporter mice and performed bulk RNA-seq to directly compare the transcriptome of the two populations. We found that the two populations showed a profoundly different transcriptional profile, with 1599 significantly differentially expressed protein coding genes (DEGs) between *Itgb8^+^* and *Itgb8^+^* CD4^+^ T_EM_ with a Ľog_2_ fold change greater than 0.5 (Figure 1e, Figure S2a-b, and Table SI). Amongst these DEGs were a number of genes known to be associated with immune suppression or regulatory function, including *Lag3, Tigit, 1110, Rnfl28* and *Pdcdl,* which showed increased expression in *Itgb8^+^* T_EM_ relative to the *Itgb8^+^* subset (Figure If). These data suggest *Itgb8^+^* CD4^+^ T_EM_ may represent a suppressive population of memory T cells. Such a function would fit with previous findings demonstrating that other immune cells capable of activating TGFβ via integrin αvβ8, including dendritic cells (DCs)^14-18^, T_REGS_ and monocytes and macrophages, are associated with immune suppressive or regulatory functions.

When sorting *Itgb8^+^* and *Itgb8^+^* CD4^+^ T_EM_ from *Itgb8* reporter mice for RNA-seq analysis, Foxp3^+^ T_REGs_ were excluded based on their expression of GFP. However, as several of the genes found to be up-regulated in *Itgb8^+^* CD4^+^ T_EM_ are also associated with regulatory immune responses, we performed a four-way bulk RNA-seq experiment to directly address whether *Itgb8^+^* Foxp3^+^ CD4^+^ T_EM_ were transcriptionally similar to CD4^+^ Foxp3^+^ T_REG_. Principle component analysis (PCA) revealed each of the four CD4^+^T cell subsets to be transcriptionally distinct from one another (Figure 1g and Table S2). Unexpectedly, we observed more variance based on *Itgb8* expression status (60%) than the difference between T_EM_ and T_REG_ (Foxp3^+^ and Foxp3^+^; 30%) (Figure 1g), a feature confirmed by correlating gene expression across groups (Figure S2c). Notably, expression of specific suppressive genes was enriched in both *Itgb8^+^* Foxp3^+^ T_EM_ and *Itgb8^+^* Foxp3^+^ T_REG_ relative to each of their respective *Itgb8^+^* counterparts (Figure 1h). However, although sharing some suppression-related gene expression, there were clear differences in expression of other T cell-associated genes (Figure S2d). Thus, these data show *Itgb8^+^* Foxp3^+^ T_EM_ are transcriptionally distinct from *Itgb8^+^* Foxp3^+^ T_REG_ and identifies a Foxp3^+^ memory T cell subset with an immune-suppressive transcriptional profile.

### *ItgbS^+^* CD4^+^ T_EM_ suppress lAV-specific CD8^+^ memory T cell responses

Next, we sought to determine the functional importance of *Itgb8* expression by CD4^+^ T_EM_ *in vivo,* in the context of an antigen-specific recall response following secondary infection. To this end, we utilised the well-characterised secondary influenza challenge model using heterologous strains of IAV. Briefly, *Itgb8^ΔCD4^* and littermate control mice were infected intranasally with the IAV strain X31 (H3N2), allowed to recover for ∼6 weeks, and then re­challenged with the IAV strain A/PR/8 (H1N1, PR8) (Figure 2a). The magnitude of the anti­viral T cell responses was tracked using major histocompatibility complex (MHC) class I- and class Il-restricted tetramers, which facilitated identification of IAV nucleoprotein (NP)^+^ specific CD8^+^ and CD4^+^ T cell responses, respectively (representative gating strategy shown in Figure S3a). We found that, in comparison to littermate controls, by day 8 post-infection *Itgb8^ΔCD4^* mice had significantly enhanced lAV-specific CD8^+^ T cell responses in the lung following re-challenge with PR8, both in terms of frequency (Figure 2b-c) and absolute cell numbers (Figure 2d), with similar increases observed in the lung-draining lymph nodes (Figure S3b-c). In contrast, lAV-specific CD4^+^ T cell responses were unaffected (Figure S3d-e). Notably, no differences in the abundance of lAV-specific CD8^+^ T cells were observed following primary infection (Figure S3f-g), indicating *Itgb8* serves to restrain cytotoxic T lymphocyte (CTL) responses following reactivation, rather than regulating their differentiation following primary infection.

**Fig. 2.**
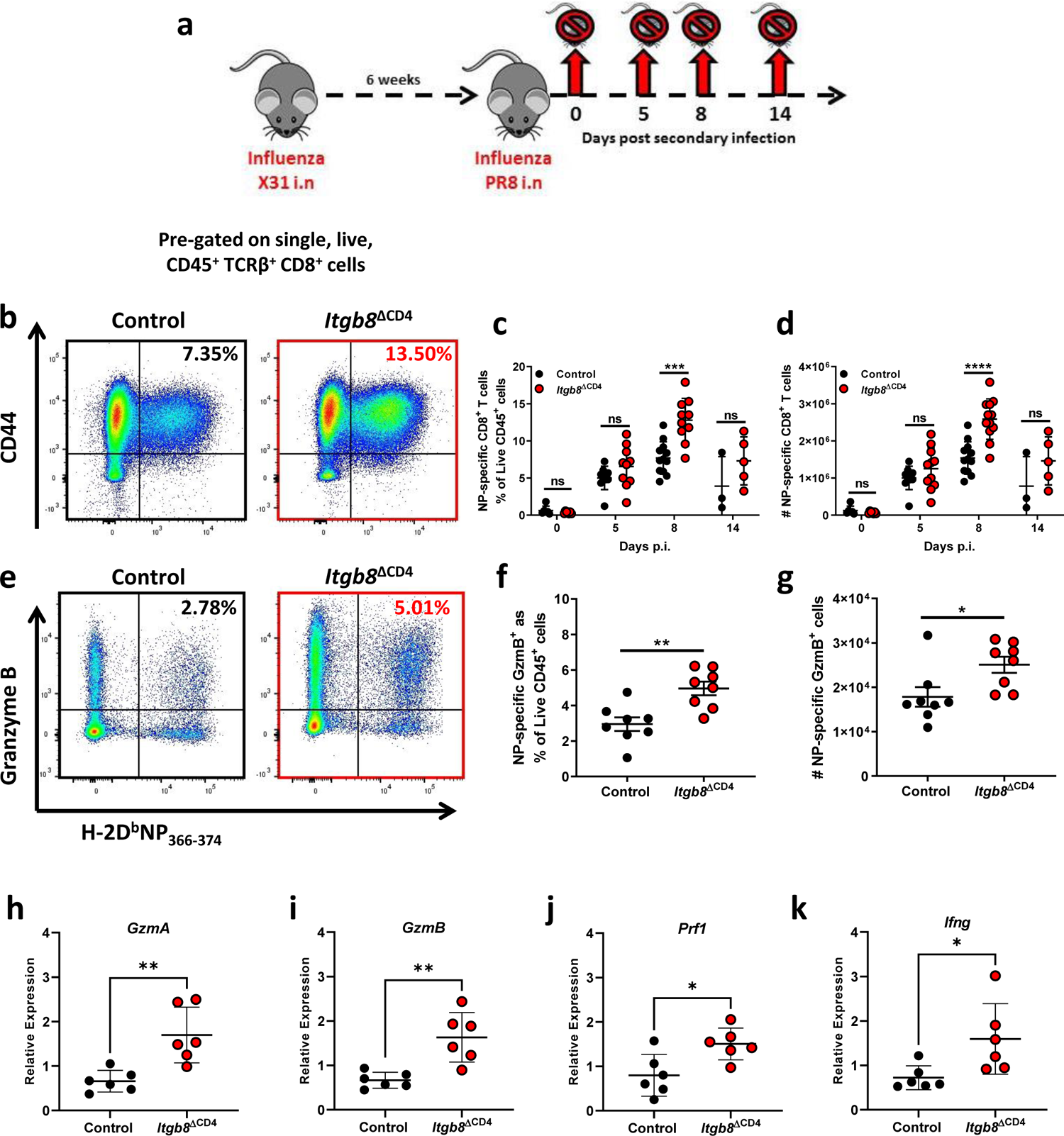
Conditional deletion of *Itgb8* on CD4+ T_EM_ enhances Influenza virus-specific CD8+ T cell responses following heterologous secondary challenge, a) Schematic of heterologous secondary Influenza A Virus challenge protocol. Littermate control and *Itgb8^ΔCD4^* mice were infected with 10^3^ PFU IAV-X31 resulting in a transient, self-resolving infection. Six weeks later, mice were re-challenged with 10^1^ PFU of the heterologous virus IAV-PR8 to facilitate assessment of lAV-specific T cell responses, b) Representative flow plots of CD44^hl^ Influenza nucleoprotein (NP[366-_3_74])-specific CD8^+^ T cells in the lungs of littermate control and *Itgb8^ΔCD4^* mice 8 days following secondary infection with IAV-PR8 as determined by tetramer staining (see Figure S3a for gating strategy). Frequencies shown are a percentage of total, live CD45^+^, single cells, c) Frequency and d) total number of NP-specific CD8^+^ T cells in the lungs of littermate control and *Itgb8^ΔCD4^* mice at indicated time-points following re­challenge with IAV-PR8. Data (n=3-10 biological replicates) from three independent experiments. Statistical significance determined by two-way ANOVA with Tukey’s multiple comparisons test ***p=≤0.001, *p=≤0.05, ns = not significant, e) Representative flow plots of total CD8^+^ T cells producing Granzyme B in the lungs of littermate control and *Itgb8^ΔCD4^* mice 5 days following secondary infection with IAV-PR8 as determined by tetramer and intracellular staining, f) Frequency (as a percentage of total live CD45^+^ cells) and g) number of Granzyme B-producing NP-specific CD8^+^ T cells in the lungs of littermate control and *Itgb8^ΔCD4^* mice 5 days following secondary infection with IAV-PR8 (n=8 biological replicates). Statistical significance determined by Mann-Whitney test **p=≤0.01, *p=≤0.05. h-k) 1 x IO^5^ NP-specific CD8^+^ T cells were FACS-purified from the lungs of littermate control and *Itgb8^ΔCD4^* mice 5 days following secondary infection with IAV-PR8 and expression of h) *Gzma,* i) *Gzmb,* j) *Prfl* and k) *Ifng* determined by RT-PCR (n=6 biological replicates from two independent experiments). Statistical significance determined by Mann-Whitney test **p=≤0.01, *p=≤0.05.

As both Foxp3^+^ T_REg_ and CD4^+^ T_EM_ express *Itgb8* (Figure 1a-c and ^12^), a caveat of the experiments using *Itgb8^ΔCD4^* mice is that expression of the integrin is ablated on both T cell subsets. Crucially however, the hyper-responsive lAV-specific CD8^+^ T cell phenotype we observed following secondary IAV infection of *Itgb8^ΔCD4^* mice is not due to a lack of *Itgb8* expression in CD4^+^ T_REG_, as deletion of *Itgb8* expression exclusively on Foxp3^+^ T_REG_ (/ŕgŵ8^flo×/flo×^ x Foxp3-Cre^12^, herein called *ltgb8^Ďfo×p3^* mice) exhibited no such phenotype after IAV re-infection (Figure S3h-i).

Next, we analysed the effector profile of lAV-specific CD8^+^ T cells in *Itgb8^ΔCD4^* mice following secondary influenza challenge. In addition to being more abundant after secondary infection, lAV-specific CD8^+^ T cells from *Itgb8^ΔCD4^* mice exhibited heightened effector properties, including enhanced production of granzyme B in the lung (Figure 2e-g) and lung­draining LN (Figure S3j-k) and increased expression of genes encoding molecules associated with cytotoxic T cell responses such as granzyme A (Gzmơ), granzyme B *(Gzmb),* perforin *(Prfl)* and interferon gamma *(Ifng)* (Figure 2h-k).

Additionally, recent evidence suggests TGFβ is crucial in the formation of CD8^+^ T_RM_, and that activation of TGFβ by DCs and epithelial cells in an αv integrin-dependent manner promotes the formation of T_RM_ at barrier sites such as the skin and intestine^20^, ^22^. However, when we analysed the development of lAV-specific CD8^+^ T_RM_ in the lung following secondary infection (see materials and methods, and Figure S3a for gating strategy), this remained intact, and was even elevated, in *Itgb8^ΔCD4^* mice (Figure S31).

Thus, taken together, these data suggest *Itgb8^+^* CD4^+^ T_EM_ have a suppressive phenotype and dampen CD8^+^ T cell numbers and effector function *in vivo* during secondary viral infection. Additionally, this cellular pathway of TGFβ activation is not involved in induction of CD8^+^T_RM_ in the lung, pointing to a dichotomy in functional outcomes depending on the cell type responsible for the activation of TGFβ.

### *ItgbS^+^* CD4^+^ T_Em_ directly suppress Influenza virus-specific CD8^+^ memory T cell responses via activation of TGFβ

Thus far, we have shown that deletion of *İtgb8* on T cells in *Itgb8^ΔCD4^* mice results in enhanced CD8^+^ T cell responses during secondary IAV infection. Furthermore, T cell expression of *Itgb8* appears to be limited to CD4^+^ T_EM_ and CD4^+^ Foxp3^+^ T_REG_ (Figure la-c and ^12^), and mice lacking expression of *Itgb8* on Foxp3^+^ T_REG_ do not show any phenotype during secondary IAV infection (Figure S3h-i). This strongly suggests that the enhanced lAV-specific CD8^+^ T cell responses observed in *Itgb8^ΔCD4^* mice is due to lack of *Itgb8* expression by CD4^+^ T_EM_. To prove this unequivocally and directly test the suppressive capacity *ofltgb8^+^* CD4^+^ T_EM_ *in vivo, we* designed an experiment to assess whether *İtgb8^+^* CD4^+^ T_EM_ were sufficient to reverse the enhanced lAV-specific CD8^+^ T cell responses in *Itgb8^ΔCD4^* mice following secondary IAV infection. To this end, we isolated *Itgb8^+^* and *Itgb8^+^* CD4^+^ T_EM_ from *Itgb8-* reporter mice following resolution of primary X31 infection and adoptively transferred them into convalescent *İtgb8^ΔCD4^* mice prior to secondary PR8 infection (Figure 3a). We found that although transfer of *Itgb8^+^* CD4^+^ T_EM_ had no effect, transfer of *Itgb8^+^* CD4^+^ T_EM_ completely reversed the enhanced NP-specific CD8^+^ T cell response to IAV re-challenge seen in *Itgb8^ΔCD4^* mice, to levels observed in littermate controls (Figure 3b-c). In addition, such an effect was not limited to NP-specific CD8^+^ T cells, as transfer of *Itgb8^+^* CD4^+^ T_EM_ also efficiently attenuated IAV polymerase (PA)-specific CD8^+^ T cell responses in *ltgb8^L·C¤Ą^* recipient mice to levels seen in littermate controls, whereas transfer of *Itgb8^+^* CD4^+^ T_EM_ had no such effect (Figure S4a-b). Thus, these data show that *Itgb8^+^* CD4^+^ T_EM_ cause suppressive effects *in vivo* and are sufficient to reverse enhanced lAV-specific CD8^+^ T cell responses in *Itgb8^ΔCD4^* mice.

**Fig. 3.**
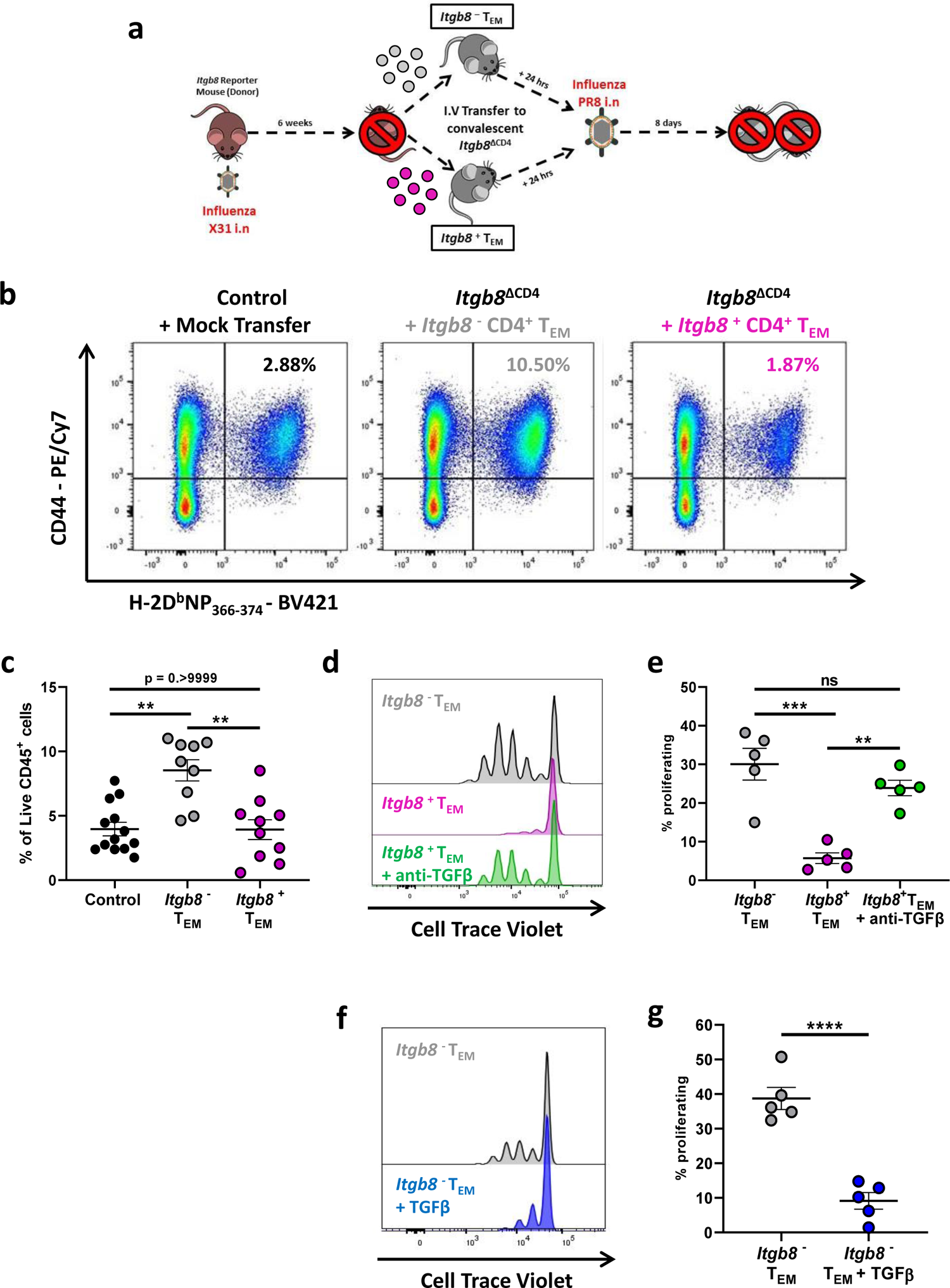
*ItgbS’* CD4^+^ T_EM_ restrain anti-viral CD8^+^ T cell responses following secondary challenge and prevent CD8^+^ T cell proliferation via TGFβ. a) Schematic of CD4^+^ T_EM_ adoptive transfer study. Littermate control, *Itgb8* and /řgðS-reporter mice were infected with 10 PFU IAV-X31. 6 weeks post-infection, *Itgb8^+^* and *Itgb8^+^* CD4^+^ T_EM_ were purified by flow cytometry from convalescent /řgðS-reporter mice and transferred intravenously into convalescent *Itgb8* recipients, 24 hours prior to heterologous re-challenge with 10 PFU IAV-PR8. b) Representative plots of NP-specific CD8^+^ T cells in the lungs of littermate control and *Itgb8^ΔCD4^* mice at day 8 post re-infection with IAV-PR8, the latter separated into two cohorts based on having received either CD4^+^ *Itgb8^+^* or *Itgb8^+^* T_EM_· c) Frequency, as a percentage of total live CD45^+^ cells, of NP-specific CD8^+^ T cells in the lungs of littermate control and *Itgb8^ΔCD4^* mice receiving either CD4^+^ *Itgb8^+^* or *Itgb8^+^* T_EM_ at day 8 post re­infection with IAV-PR8. Data (n=≥9) from three independent experiments. Statistical significance determined by Kruskall-Wallis with Dunn’s multiple comparisons test **p=≤0.01, ns = not significant, d) Representative histograms of Cell Tracer Violet-labelled naïve CD8^+^ T cell, stimulated with anti-CD3 and anti-CD28 antibodies and co-cultured in a 1:1 ratio with *Itgb8^+^* or *Itgb8^+^* CD4^+^ T_EM_ for 72 hours. Grey peaks represent CD8^+^ T cells co­cultured with *Itgb8^+^* CD4^+^ T_EM_, purple peaks signify CD8^+^ T cells co-cultured with *Itgb8^+^* CD4^+^ T_EM_ and green peaks represent CD8^+^ T cells co-cultured with *Itgb8^+^* CD4^+^ T_EM_ in the presence of a TGFβ blocking antibody, e) Pooled data for frequency of CD8^+^ T cells undergoing proliferation under each condition in d (n=5 biological replicates from two independent experiments). Statistical significance determined by one-way ANOVA with Tukey’s multiple comparisons test ***p=≤0.001, **p=≤0.01, ns = not significant, f) Representative histograms and g) collated data of naïve CD8^+^ T cell proliferative responses following co-culture in a 1:1 ratio with *Itgb8^+^* CD4^+^ T_EM_ plus or minus recombinant TGFβ for 72 hours. Grey peaks/dots represent CD8^+^ T cells co-cultured with *Itgb8^+^* CD4^+^ T_EM_, blue plots/dots represent CD8^+^ T cells co-cultured 1:1 with *Itgb8^+^* CD4^+^ T_EM_ in the presence of recombinant TGFβ (n=5 biological replicates from two independent experiments). Statistical significance determined by unpaired student’s t-test ****p=≤0.0001.

In light of these findings, we next sought to elucidate the mechanism(s) by which *İtgb8^+^* CD4^+^ T_EM_ can suppress CD8^+^ T cell responses after viral re-infection. As expression of *Itgb8* by CD4^+^ T_EM_ enables them to activate TGFβ (Figure Id), and TGFβ can directly inhibit CD8^+^ T cells^23^, we postulated that *Itgb8^+^* CD4^+^ T_EM_ could directly suppress CD8^+^ T cell responses via their ability to activate TGFβ. To investigate this hypothesis, we performed an *ex vivo* CD8^+^ T cell proliferation assay and tested the ability of *Itgb8^+^* and *Itgb8^+^* CD4^+^ T_EM_ to modulate the proliferative response. To this end, we isolated *Itgb8^+^* and *Itgb8^+^* CD4^+^ T_EM_ from *Itgb8* reporter mice and co-cultured each of these subsets separately with Cell Trace Violet (CTV)-labelled CD8^+^ T cells that were activated with anti-CD3ξ and anti-CD28 antibodies. We found that CD8^+^ T cell proliferation was significantly inhibited when responder cells were co-cultured with *Itgb8^+^,* but not *Itgb8^+^* CD4^+^ T_EM_ (Figure 3d-e). Furthermore, the inhibition of CD8^+^ T cell proliferation by *Itgb8^+^* CD4^+^ T_EM_ was wholly dependent on active TGFβ, as this inhibition was completely reversed when the cells were co-cultured in the presence of a TGFβ blocking antibody (Figure 3d-e). Moreover, addition of exogenous, active TGFβ to co-cultures of *Itgb8^+^* CD4^+^ T_EM_ with CD8^+^ responders also significantly reduced CD8^+^ T cell proliferation (Figure 3f-g), further implicating TGFβ as a key factor involved in immune suppression mediated by *Itgb8^+^* CD4^+^ T_EM_.

Taken together, these data show that *Itgb8^+^* CD4^+^ T_EM_ restrain lAV-specific CD8^+^ T cells *in vivo,* a function facilitated via their ability to activate TGFβ in an integrin αvβδ-dependent manner, which can act directly on CD8^+^ T cells to inhibit their proliferation.

### *ItgbS^+^* CD4^+^ T_EM_ slow viral clearance but are vital in preventing lung pathology following viral re-infection

We next sought to determine the physiological relevance of *Itgb8^+^* CD4^+^ T_EM_-mediated control of anti-viral memory T cell responses. To this end, we first analysed the ability of *Itgb8^ΔCD4^* mice and their littermate controls to clear the virus after secondary IAV infection. We found that *Itgb8^ΔCD4^* mice showed significantly more robust expulsion of the virus, with reduced copies of viral RNA detected at days 5 and 8 post-secondary infection compared to littermate controls (Figure 4a). In contrast, we did not observe any difference in the ability of *Itgb8^ΔCD43^* mice to clear viral load after secondary infection compared to littermate controls (Figure S5a), providing strong evidence that the lack of *Itgb8* expression on Foxp3^+^ T_REG_ is not responsible for the enhanced viral clearance in *Itgb8^ΔCD4^* mice. Taken together, our data indicate that *Itgb8^+^* CD4^+^ T_EM_-mediated restraint of CD8^+^ T cells during secondary IAV infection limits the ability of mice to rapidly clear the virus.

**Fig. 4.**
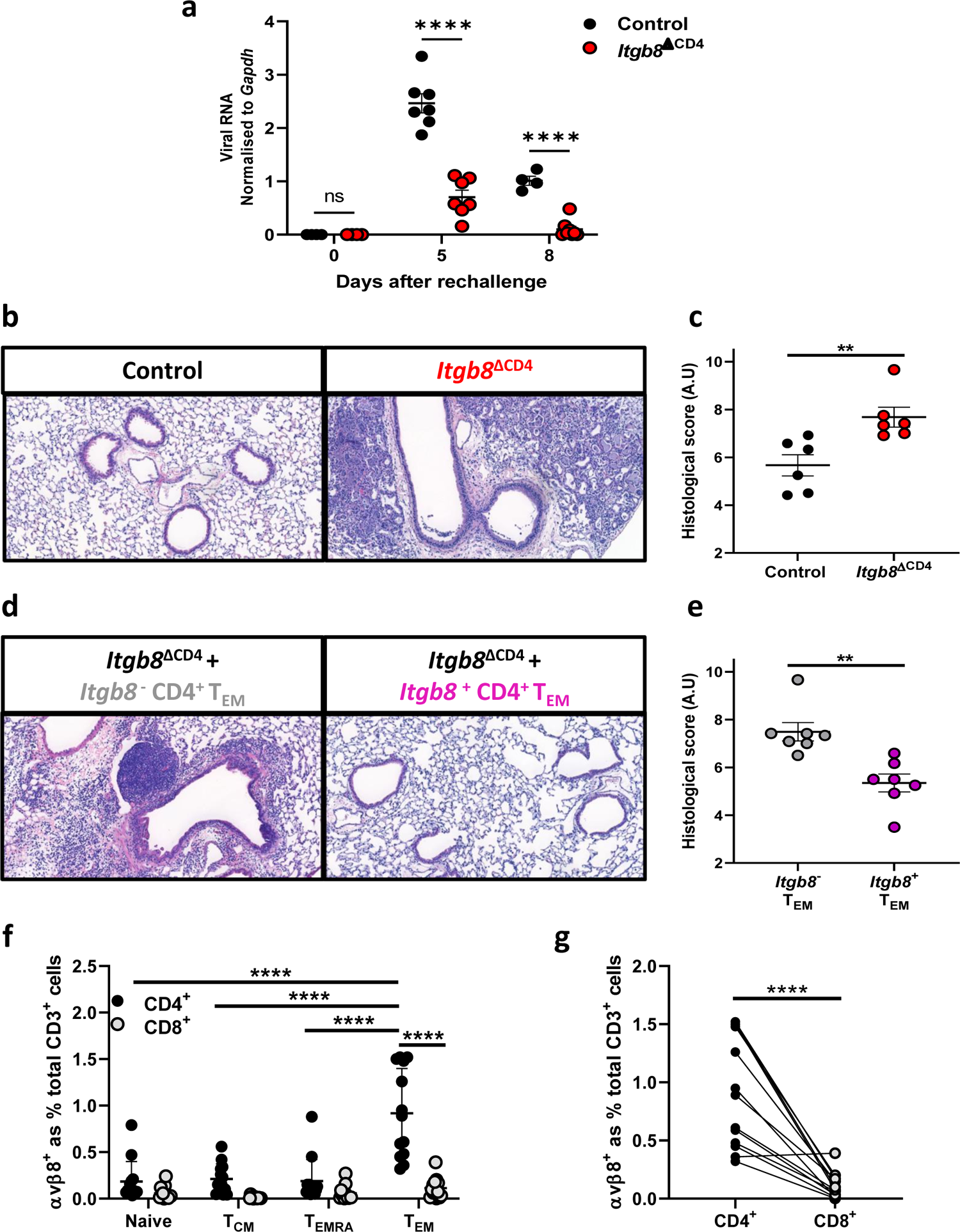
Conditional deletion of *Itgbδ* on CD4+ T_EM_ facilitates rapid viral clearance but induces significant tissue pathology, a) Viral Matrix (M) protein mRNA in the lungs of littermate control and *Itgb8^ΔCD4^* mice at indicated time-points following re-challenge with IAV-PR8 as determined by RT-PCR (n=≥4 biological replicates per-timepoint from two independent experiments). Statistical significance determined by two-way ANOVA with Šídák’s multiple comparisons test ****p=≤0.0001. b) Representative images of H&E-stained lung tissue sections from littermate control and *Itgb8^ΔCD4^* mice 8 days following re-challenge with IAV-PR8. c) Inflammation scores of blind-scored H&E-stained lung tissue sections from littermate control and *Itgb8^ΔCD4^* mice 8 days following re-challenge with IAV-PR8 (n=6 biological replicates). Statistical significance determined by Mann-Whitney test **p=≤0.01. d) Representative images of H&E-stained lung tissue sections from *Itgb8^ΔCD4^* mice 8 days following re-challenge with IAV-PR8. 24 hours prior to re-challenge, *Itgb8^ΔCD4^* mice were adoptively transferred with either *Itgb8^+^* CD4^+^ T_EM_ (grey) or *Itgb8^+^* CD4^+^ T_EM_ (purple) purified from convalescent ItgbS-reporter mice, e) Inflammation scores of blind-scored H&E-stained lung tissue sections from *Itgb8^ΔCD4^* recipient mice 8 days following re-challenge with IAV-PR8. Statistical significance determined by Mann-Whitney test **p=≤0.01. f) Frequency of integrin αvβ8-expressing cells amongst CD4^+^ and CD8^+^ T cell subsets in human PBMCs. Frequencies shown are a percentage of total CD3^+^ cells (n=10 individual donors). Statistical significance determined by two-way ANOVA with Tukey’s multiple comparisons test, g) Pairwise comparison of integrin αvβ8-expressing CD4^+^ and CD8^+^ T_Cm_ in human PBMCs. Statistical significance determined by two-tailed, paired student’s t-test ****p=≤0.0001.

Our studies demonstrated a novel role for *Itgb8*-expressing CD4^+^ T_EM_ in restraining anti-viral CTL responses and slowing clearance of the virus. We next wanted to determine the functional importance of a pathway that essentially limits host immunity to pathogenic challenge. Based on the potent cytotoxic profile of lAV-specific CD8^+^ T cells in *itgb8^ΔCD4^* mice (Figure 2e-k), we hypothesised that this pathway may be important in limiting infection-induced inflammation and lung pathology. Indeed, *itgb8^ΔCD4^* mice exhibited significantly enhanced lung inflammation compared to littermate control animals after secondary infection, with inflammatory cells locating around airways and blood vessels and infiltrating the parenchymal area (Figure 4b-c). There was no difference in lung pathology between *Itgb8^ΔCD4^* mice and littermate controls after primary IAV infection (Figure S5b-c), again indicating a role for *Itgb8^+^* CD4^+^ T_EM_ in controlling T cell responses during re-infection rather than primary infection.

We next determined whether enhanced pathology observed after secondary IAV infection in *Itgb8*^ΔCD4^ mice was directly due to the absence of *Itgb8^+^* CD4^+^ T_EM_. T_o_ this end, we analysed lung pathology following secondary IAV infection of *Itgb8^ΔCD4^* mice that received either *Itgb8^+^* or *Itgb8^+^* CD4^+^ T_EM_. Transfer of *Itgb8^+^* CD4^+^ T_EM_, but not *Itgb8^+^* CD4^+^ T_EM_, completely reversed the enhanced lung pathology seen in *Itgb8^ΔCD4^* mice after secondary infection (Figure 4d-e), restoring histology and inflammation scores to that seen in control mice (Figure 4b-c). Taken together, these data strongly indicate that although *Itgb8^+^* CD4^+^ T_EM_ slow the ability of the host to clear viral load, they are essential in restraining pathogenic anti-viral CD8^+^ T cell responses to prevent tissue pathology following re­infection.

Having discovered that integrin αvβ8 expression identifies a sub-population of immune suppressive murine CD4^+^T_EM_, which can restrain effector recall responses and prevent overt inflammatory sequelae, we sought to determine whether a similar population might be present in humans. To test this, we isolated human peripheral blood mononuclear cells (PBMCs) and assessed expression of integrin αvβ8 on human T cell subsets by flow cytometry using a monoclonal antibody specific for human αvβ8^15^ (gating strategy shown in Figure S6). In agreement with our pre-clinical data, expression of integrin αvβ8 was restricted primarily to CD4^+^ T_Cm_ in human PBMCs (Figure 4f-g).

Together, our data for the first time show that a subset of cells classed as CD4^+^ memory T cells can be suppressive during re-infection. This suppressive function is reliant on their expression of an integrin, αvβ8, which enables them to activate TGFβ. We find that these cells are crucial in restraining CD8^+^T cell responses during re-infection with IAV in the lung but are also present in the intestine. Thus, important future work will determine if these cells have a broader function in regulating secondary immunity in other organs to a wide range of challenges, or if the cells are specific to pulmonary immunity.

Our findings have important implications to how the immune system balances responses to re-infection. Although immunological memory has evolved to promote more robust responses to subsequent infections, it now appears that memory T cell subsets can also limit the robustness of this response. In situations where re-infection is proving hard to irradicate, it could be beneficial to inhibit the function of these suppressive memory T cells, via blocking integrin αvβ8 and therefore the ability of these cells to activate TGFβ, to boost CD8^+^ T cell responses.

However, caution would be required with any such approaches, as these suppressive memory T cells appear critical in limiting host pathology during re-infection. Thus, strategies would have to balance the beneficial effects of promoting immunity to re-infection, with potential tissue damage associated with an exacerbated response. A phase I clinical trial is already underway in cancer using an antagonist of integrin αvβ8 to block TGFβ activation, based on successful pre-clinical studies^24^, ^26^. Based on our findings that this molecular pathway is important in limiting tissue pathology to re-infection, attention to the infection status of the patients during treatment periods may be prudent.

## Materials and methods

### Mice

Conditional deletion of *Itgb8* (β8 integrin) expression specifically in T cells or T_REg_ was achieved by crossing mice with a conditional floxed allele of *Itgb8* with CD4-Cre^14^ or Foxp3-YFP-Cre^12^ mice. All mice were used for experiments at 8 to 12 weeks of age, with littermates used as control. Itgb8-IRES-TdTomato mice, generated as previously described^13^, were provided by Dr. H. Paidassi and bred at the University of Manchester. All experiments were approved under a project license granted by the Home Office UK, reviewed by the University of Manchester Animal Welfare and Ethical Review Body and were performed in accordance with the UK Animals (Scientific Procedures) Act of 1986. Mice were bred and maintained in specific pathogen-free conditions at Bio Safety Level 2 with a 12-hour light/dark cycle and food and water *ab libitum*.

### Influenza A Virus Infections

*Itgb8* conditional knockout mice and their littermate controls were anaesthetised with 2.5% inhalable isoflurane and inoculated with 10^3^ PFU of the IAV strain X31 (H3N2) or 10^1^ PFU of the IAV strain A/PR/8 (H1N1, PR8) intranasally, resulting in a transient infection and mild to moderate weight loss. For secondary IAV challenge studies mice were inoculated with 10^3^ PFU IAV-X31 and subsequently re-challenged ∼6 weeks later with 10^1^ PFU of the heterologous strain IAV-PR8 and immune responses assessed at time-points indicated in the figure legends.

#### Lung digest

Lungs were transferred into 7 ml bijoux tubes containing 500 µl digestion mix consisting of 5 mg/ml collagenase I (Gibco) and 10 µg/ml DNase I (Sigma) in RPMI and finely minced with scissors. An additional 1.5 ml of digestion mix was then added to samples and incubated in a shaking incubator at 37°C for 40 minutes. Enzymes were quenched with ice-cold 10% RPMI medium containing 10% FCS, and cells pelleted via centrifugation at 400 x g for 8 minutes. Cell pellets were re-suspended in 2 ml red blood cell lysis buffer (Sigma) and washed with ice-cold 10% FCS in RPMI prior to staining for flow cytometry.

#### Flow Cytometry

Cell suspensions were transferred into 5 ml round bottom polystyrene tubes, washed in PBS and stained with fixable viability Zombie dyes (Biolegend) diluted in PBS. Staining was quenched by washing in FACS buffer (PBS with 2 mM EDTA and 2% FCS) after which cells were incubated with anti-CD16/32 (Fc block, clone 2.4G2, BD Biosciences) for 15 minutes at 4°C to prevent non-specific binding. Without washing, antibodies specific for surface markers were diluted in FACS buffer and added 1:1 directly onto samples and incubated for a further 20 minutes at 4°C protected from light. Stained samples were then washed in PBS and either re-suspended in FACS buffer for immediate sorting on a BD FACSAria™ cell sorter or fixed using eBioscience™ Foxp3 / Transcription Factor Staining Buffer Set as per manufacturer’s guidelines. For experiments requiring T cell sorting, single cell suspensions underwent a negative selection pre-enrichment using EasySep™ pan-T cell or memory CD4^+^ T cell isolation kits as per manufacturer’s guidelines (Stemcell™ Technologies).

To stain for intracellular markers or cytokines, fixed/permeabilsed samples were incubated with antibodies diluted in IX permeabilisation buffer. Samples were washed twice and re­suspended in FACS-EDTA prior to acquisition on a BD LSR Fortessa™ or BD Symphony™. Antibodies used for flow cytometry and FACS were purchased from Biolegend and/or eBioscience™ and are listed in Supplementary Table 3.

To assess T_RM_ responses, intra-vascular labelling was performed as previously described^27^. Briefly, 3 µg anti-CD45.2 (104) conjugated to either FITC or APC was administered via i.v injection into the tail vein in a final volume of 200 µl 3-5 minutes prior to performing euthanasia. All flow cytometry data was analysed using FlowJo V10 software (Tree Star).

#### Tetramer Staining

For assessment of Influenza virus-specific T cell responses, cells were transferred into 5 ml round bottom polystyrene tubes, washed and incubated with Fc block for 15 minutes at room temperature to prevent non-specific binding. Tetramers specific for CD8^+^ T cells (H-2D(b) Influenza A NP_366___374_ (ASNENMETM) and H-2D(b) Influenza A PA_224_-_233_ (SSLENFRAYV)) and CD4^+^ cells (l­A(b) Influenza A NP_3_ıı-3_25_ (QVYSLIRPNENPAHK)) recognising immunodominant epitopes were obtained from the NIH tetramer facility at Emory University and added at 5 µg/ml in RPMI and incubated for a minimum of 1 hour at 37°C with regular agitation. Cells were washed in PBS and staining performed as described below thereafter.

#### Quantitative PCR

FACS-purified cells were sorted into complete media, washed in sterile PBS and re­suspended in RLT buffer (Qiagen) containing 1% β-mercaptoethanol. For assessment of viral mRNA in lung tissue a single lobe from the right lung was collected into a sterile 1.5 ml tube containing RNA later and subsequently transferred into a 2ml tube containing RLT buffer and a single 2 mm stainless steel bead. Tissue was homogenised into suspension using a Qiagen Tissue Lyser. RNA was extracted from both cells and lysed tissue samples using Qiagen RNeasy Mini Kit as per manufacturer’s instructions. Reverse transcription of RNA to cDNA was performed using the High Capacity RNA-to-cDNA Kit (Thermo Fisher Scientific) and gene expression was assayed using PowerUp™ SYBR™ Green Master Mix (Applied Biosystems). Reactions were run on a real-time PCR system (ABI 7500; Applied Biosystems). Data for samples were normalized to *Gapdh* and are shown as a fold change compared with the controls.

#### TGF-β Activation Assay

FACS-purified T cell subsets were incubated overnight with a TGF-β reporter cell line^28^ and luciferase activity detected via the Luciferase Assay System (Promega). TGF-β activity was determined as previously described^16^.

#### RNA sequencing (RNA-seq)

Total RNA was submitted to the Genomic Technologies Core Facility (GTCF) at the University of Manchester and the quality and integrity of the RNA samples assessed using a 4200 TapeStation (Agilent Technologies) prior to the generation of libraries using the lllumina^®^ Stranded mRNA Prep. Ligation kit (lllumina, Inc.) according to the manufacturer’s protocol. Briefly, total RNA was used as input material from which polyadenylated mRNA was purified using poly-T, oligo-attached, magnetic beads. Next, the mRNA was fragmented under elevated temperature and then reverse transcribed into first strand cDNA using random hexamer primers and in the presence of Actinomycin D (thus improving strand specificity whilst mitigating spurious DNA-dependent synthesis). Following removal of the template RNA, second strand cDNA was then synthesized to yield blunt-ended, double-stranded cDNA fragments. Strand specificity was maintained by the incorporation of deoxyuridine triphosphate (dUTP) in place of dTTP to quench the second strand during subsequent amplification.

Following a single adenine (A) base addition, adapters with a corresponding, complementary thymine (T) overhang were ligated to the cDNA fragments. Pre-index anchors were then ligated to the ends of the double-stranded cDNA fragments to prepare them for dual indexing. A subsequent PCR amplification step was then used to add the index adapter sequences to create the final cDNA library. The adapter indices enabled the multiplexing of the libraries, which were pooled prior to cluster generation using a cBot instrument. The loaded flow-cell was then paired-end sequenced (76 + 76 cycles, plus indices) on an lllumina HiSeq4000 instrument. Finally, the output data was demultiplexed and BCL-to-Fastq conversion performed using lllumina’s bcl2fastq software (v2.20.0.422).

#### RNA-seq data analysis

The quality of the stranded paired-end RNA-seq reads were assessed using FastQC (vθ.11.3; https://www.bioinformatics.babraham.ac.uk/projects/fastqc/) and FastQ Screen (vO.13.O; https://www.bioinformatics.babraham.ac.uk/projects/fastq_screen/), followed by adapter removal and low-quality bases and reads trimming using BBDuk (BBMap suite v36.32; http://sourceforge.net/projects/bbmap/). Processed reads were mapped against the mouse reference genome (mmlO) and gene annotation from Gencode (vM24) using STAR (v2.7.2b; https://doi.org/10.1093/bioinformatics/bts635). The “--quantMode GeneCounts” option was used to obtain read counts per gene from STAR.

Differential gene expression analysis was performed in the R environment using the Bioconductor package DESeq2 (vl.30.1; https://doi.org/10.1186/sl3059-014-0550-8). Genes were considered significantly differentially expressed with a false discovery rate cut­off of 0.05. The lfcShrink function (with apeglm method) was applied to produce a more accurate Iog2 fold change estimates. The counts function (with normalized = TRUE) was used to produce normalised counts, where it uses the median-of-ratios method to determine scaling factors. Data transformation was performed using the rlog and vst functions to obtain regularised logarithm and variance stabilizing transformation expression values respectively, rlog transformed values were used to produce principal component analysis plots using the prcomp function from the R package stats, vst values were used to produce sample-to-sample correlation using the cor function from the R package stats. Heatmaps and correlation plots were visualised using the R package pheatmap (vl.0.12).

#### T cell co-culture assays

Spleens from Itgb8-IRES-TdTomato mice were manually dissociated, filtered through 40 µm cell strainers and T cells enriched by negative selection using an EasySep™ pan-T cell enrichment kit (Stemcell Technologies) as per manufacturer’s guidelines. Splenic CD4^+^ T_Cm_ and responder CD8^+^ T cells from Itgb8-IRES-TdTomato mice were stained with cell surface markers and further purified by FACS using a BD FACSAria™. CD8^+^ T cells were subsequently stained with CellTrace™ Violet proliferation dye (Thermo Fisher Scientific) and plated in a 1:1 ratio with *Itgb8^+^* or *Itgb8^+^* CD4^+^ T_Cm_ in RPMI containing 0.88 mM L-Glutamine (Gibco), 0.88 mM sodium pyruvate (Gibco), 0.04 mM β-mercaptoethanol, 0.88% (v/v) MEM non­essential amino acids (Gibco), 0.35% (v/v) 100X MEM Vitamins (Gibco), 4.4 U/ml penicillin (Gibco), 4.4 µg/ml streptomycin (Gibco) and 10% (v/v) heat-inactivated fetal calf serum (FCS) (Sigma). Cells were added to sterile 96 well U-bottom tissue culture plates pre-coated with purified NA/LE anti-CD3 (Clone 145-2C11) and soluble NA/LE anti-CD28 (Clone 37.51; both BD Biosciences) added to provide co-stimulation. CD8^+^ T cell proliferative responses were measured after 72 hours by assessing dye dilution across cell divisions on a BD Symphony™.

#### Adoptive transfer studies

For adoptive transfer studies, *Itgb8^+^* or *Itgb8^+^* CD4^+^ T_EM_ were FACS-purified from the spleens of convalescent Itgb8-IRES-TdTomato mice and 2 x 10^5^ of either cell type administered i.v into the tail vein of IAV-X31 convalescent control or *Itgb8^ΔCD4^* mice 1 day prior to re­-challenge with IAV-PR8.

#### Histological Assessment of Inflammation

Mice were euthanised via intra-peritoneal injection of pentobarbitone and the thoracic cavity opened to expose the lungs and trachea. The right lobes of the lungs were tied off and removed for assessment of immune cell populations by flow cytometry. The remaining left lobe was subsequently perfused with ∼10 ml 1 mM EDTA in ice-cold PBS to remove blood/circulating cells. The internal structure of the left lobe was then fixed by inflating with 1 ml 10% formalin via the trachea and the entire lobe subsequently immersed in 10% formalin prior to transfer into 70% ethanol after 24 hours. Fixed lungs were embedded in paraffin and sliced into 5 µm sections prior to staining with hematoxylin and eosin. Histopathology was scored at 10x magnification of a light microscope according to a set of custom-designed criteria to assess inflammation around airways, blood vessels and parenchyma as previously described^29, 30^. Six individual snapshots of each tissue section were analysed in a blinded manner by two independent researchers and results collated.

#### Statistical Analysis

Results are expressed as mean ± SEM. All statistical analysis was performed using GraphPad Prism software (v9.4.1) and tests used are stated in the figure legends.

## Supplementary Figure Legends

**Fig. SI *Itgb8*-expressing CD4* T_EM_ are present in non-lymphoid tissues,** a) Representative gating strategy for analysing and sorting murine splenic T cell subsets, b) Representative flow plots and c) collated data showing *Itgb8* expression in lung CD4^+^ T cell subsets of *Itgb8-*reporter mice, d) Representative flow plots and c) collated data showing *Itgb8* expression in small intestinal CD4^+^ T cell subsets of ItgbS-reporter mice. Data presented as frequency of *Itgb8^+^* cells as a percentage of each parent population (n=3 biological replicates for each tissue, representative of two independent experiments). Statistical significance determined by two-way ANOVA with Tukey’s multiple comparisons test ****p=≤0.0001, ns = not significant.

**Fig. S2 *Itgb8*-expressing CD4^+^ T_EM_ and T_REG_ possess a characteristic immune suppressive transcriptional profile,** a) Heatmap of Z-scores of regularized variance stabilizing transformed read counts from bulk RNA-seq (n=6) analysing CD4^+^ *Itgb8^+^* T_EM_ (left, black) and CD4^+^ *Itgb8^+^* T_EM_ (right, red), with differential (Ľog_2_fold change > 0.5), statistically significant (adjusted p < 0.05) genes indicated, b) Principal component analysis (PCA) of bulk RNA-seq experiment performed on flow cytometry-purified splenic *Itgb8^+^* T_EM_ versus *Itgb8^+^* T_EM_ (n = 6). c) Sample similarity heatmap showing Pearson correlation between samples calculated using regularized logarithmic (rlog)­transformed read counts, d) Heatmap of Z-scores of regularized variance stabilizing transformed read counts of selected genes associated with T cell biology.

**Fig. S3 Analysis of Influenza virus-specific T cell responses following primary and heterologous secondary challenge in *Itgb8^åc¤íi^* and /*Itgb8*^ΔFoxp3^ mice,** a) Representative gating strategy used to identify Influenza NP-specific CD8^+^ T cells, as well as tissue-resident cells, in the lung of mice, b) Representative flow plots and c) collated data for the frequency of CD44^hl^ Influenza NP-specific CD8^+^ T cells in the draining mediastinal lymph nodes of littermate control and *Itgb8^ΔCD4^* mice at indicated time-points post-secondary IAV-PR8 infection (n= ≥6 biological replicates from two independent experiments). Frequencies shown are as a percentage of total CD8^+^ T cells per node. Statistical significance determined by two-way ANOVA with Tukey’s multiple comparisons test **p=≤0.01. d) Representative flow plots and e) collated data for the frequency of CD44^hl^ Influenza NP-specific CD4^+^ T cells in the lungs of littermate control and *ltgb8^L·C¤Ą^* mice at indicated time-points post re­challenge infection with IAV-PR8 (n=≥4 biological replicates per timepoint, from three independent experiments). Frequencies shown are as a percentage of total, live CD45^+^, single cells per lung. Statistical significance determined by two-way ANOVA with Tukey’s multiple comparisons test, f) Representative flow cytometry plots and g) collated data for the frequency of NP-specific CD8^+^T cells in the lungs of littermate control and *Itgb8^ΔCD4^* mice at indicated time-points post primary infection with IAV-PR8 (n=≥3 biological replicates per timepoint representative of three independent experiments). Statistical significance determined by two-way ANOVA with Tukey’s multiple comparisons test, h) Representative flow cytometry plots and i) collated data for the frequency of CD44^hl^ Influenza NP-specific CD8^+^ T cells in the lungs of littermate control and *Itgb8^ΔCD4^* mice at indicated time-points following secondary infection with IAV-PR8 (n=10 biological replicates from two independent experiments, n=3 for day 0 controls). Statistical significance determined by two-way ANOVA with Tukey’s multiple comparisons test, j) Representative flow cytometry plots and k) collated data for frequency of Granzyme B-producing NP-specific CD8^+^ T cells in the mediastinal lymph nodes of littermate control and *Itgb8^+^* mice at indicated time-points post-secondary infection with IAV-PR8. Statistical significance determined by two­way ANOVA with Tukey’s multiple comparisons test **p=≤0.01. (n=≥4 biological replicates from two independent experiments). Frequencies shown are as a percentage of total CD8^+^ T cells per node. I) Frequency of intra-vascular (CD45.2 I.V+) and extra-vasculature (CD45.2-) NP-specific CD8^+^ T cells in the lungs of littermate control and *Itgb8^+^* mice 8 days following secondary infection with IAV-PR8, as determined by intra-vascular labelling and tetramer staining (n=5 biological replicates from two independent experiments). Statistical significance determined by two-way ANOVA with Tukey’s multiple comparisons test ****p=≤0.0001, *p=≤0.05.

**Fig. S4 *ItgbS^+^* CD4^+^ T_EM_ restrain anti-lAV PA-specific CD8^+^ T cell responses following secondary challenge,** a) Representative flow cytometry plots of viral polymerase (PA_[2_24-233])-specific CD8^+^ T cells in the lungs of littermate control and *Itgb8^ΔCD4^* mice at day 8 post re­infection with IAV-PR8, the latter separated into two cohorts on the basis of having received *Itgb8^+^* or *Itgb8^+^* CD4^+^ T_EM_ intravenously one day prior to re-challenge, b) Frequency, as a percentage of total, live CD45^+^ single cells, of PA-specific CD8^+^ T cells in the lungs of littermate control and both cohorts of*Itgb8^ΔCD4^* mice at day 8 post re-infection with IAV-PR8 (n=≥9 biological replicates from three independent experiments). Statistical significance determined by Kruskall-Wallis with Dunn’s multiple comparisons test *p=≤0.05, ns = not significant.

**Fig. S5 Expression of *ItgbS’* on Foxp3^+^ T_reg_ does not control viral titre and *ItgbS’* CD4^+^ T_EM_ do not control lung pathology following primary IAV challenge,** a) Viral Matrix (M) protein mRNA in the lungs of littermate control and *Itgbδ^+¤×p3^* mice at indicated time-points following heterologous re-challenge with IAV-PR8 as determined by RT-PCR (n=3-5 biological replicates per-timepoint). Statistical significance determined by two-way ANOVA with Tukey’s multiple comparisons test, b) Representative images of H&E-stained lung tissue sections from littermate control and *Itgb8^ΔCD4^* mice 6 weeks following primary infection with IAV-PR8. c) Inflammation scores of blind-scored H&E-stained lung tissue sections from littermate control and *Itgb8^+^* mice 6 weeks following primary infection with IAV-PR8 (n=4 biological replicates). Statistical significance determined by Mann-Whitney test.

**Fig. S6 Gating strategy for assessment of integrin αvβ8 expression of human T cell subsets.** Representative gating strategy used to identify naïve T cells, T_CM_, T_EM_ and T_EMra_ in human PBMC samples isolated from whole blood by Ficoll density gradient centrifugation.

**Table SI: Differentially expressed genes between *ItgbS’* and *ItgbS* CD4^+^T_EM_.** List of 1599 differentially expressed protein-encoding genes between *Itgb8^+^* and *Itgb8^+^* CD4^+^ T_EM_ with a Ľog_2_ fold change greater than 0.5 and adjusted p-value of <0.05 (from RNA-seq experiment shown in Figure le and Figure S2a-b).

**Table S2: Differentially expressed genes between *Itgbδ\** and *Itgbδ’* CD4+ T_EM_ and T_REG_. List** of differentially expressed protein-encoding genes from all possible two-way comparisons between *Itgb8^+^* and *Itgb8^+^* CD4+ T_EM_ and T_REG_, where a gene displayed a Log2 fold change greater than 0.5 and adjusted p-value of <0.05 for any comparison (from RNA-seq experiment shown in Figure lg and Figure S2c-d).

**Table S3: List of flow cytometry antibodies and reagents used.**

