## Supplementary Figures for "A subset of CD4+ effector memory T cells limit immunity to pulmonary viral infection and prevent tissue pathology via activation of latent TGFβ"

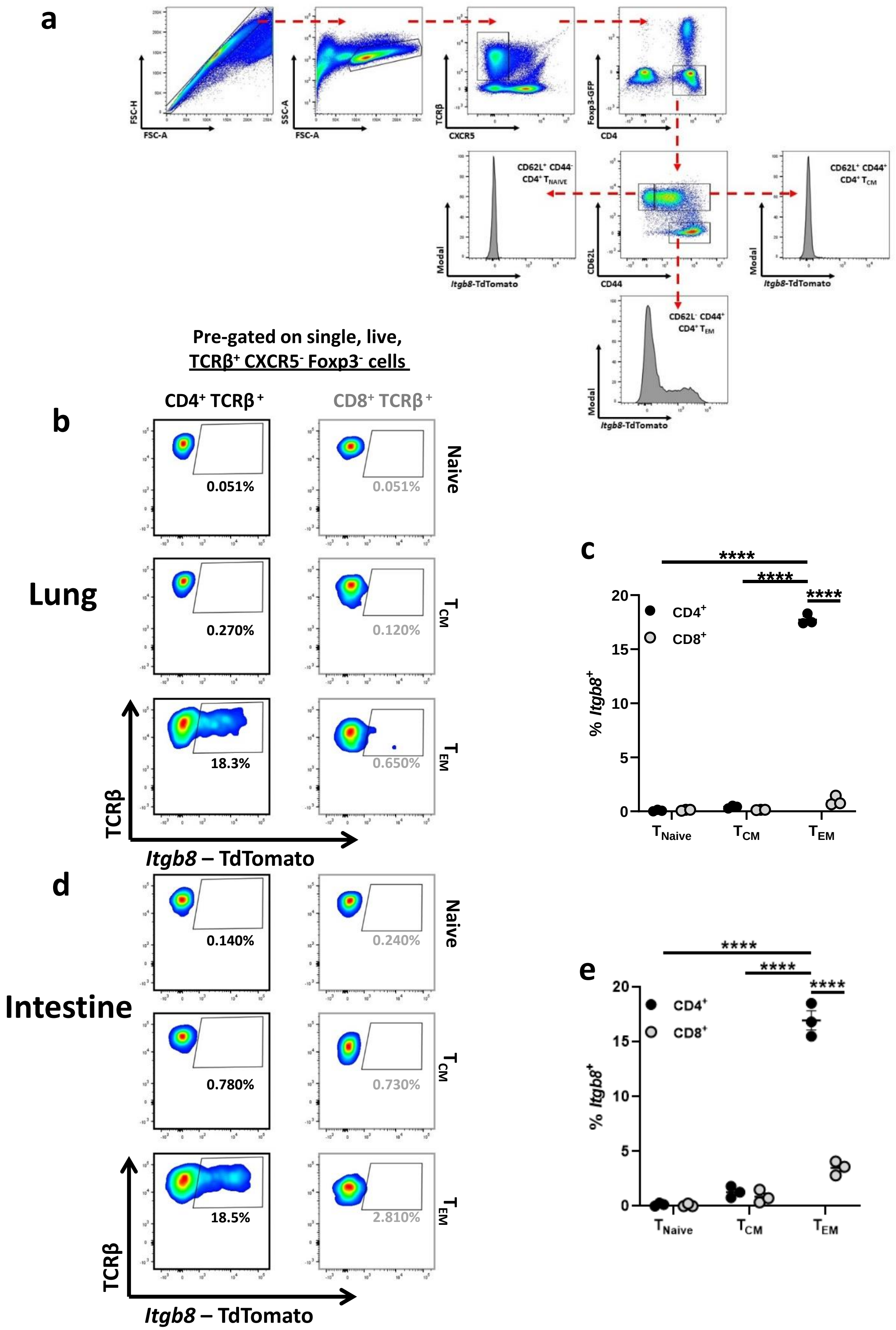


### Supplementary Figure 1

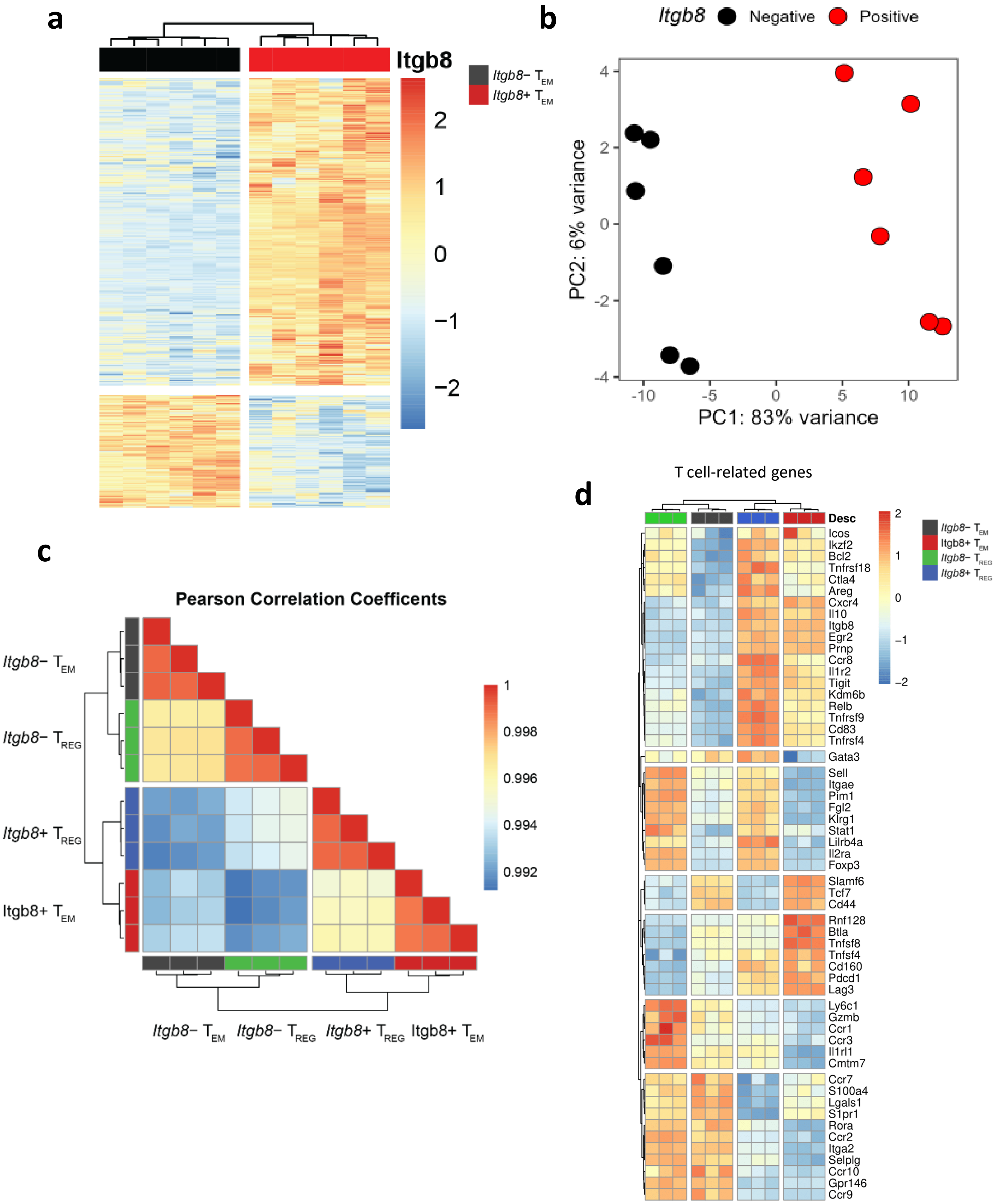


### Supplementary Figure 2

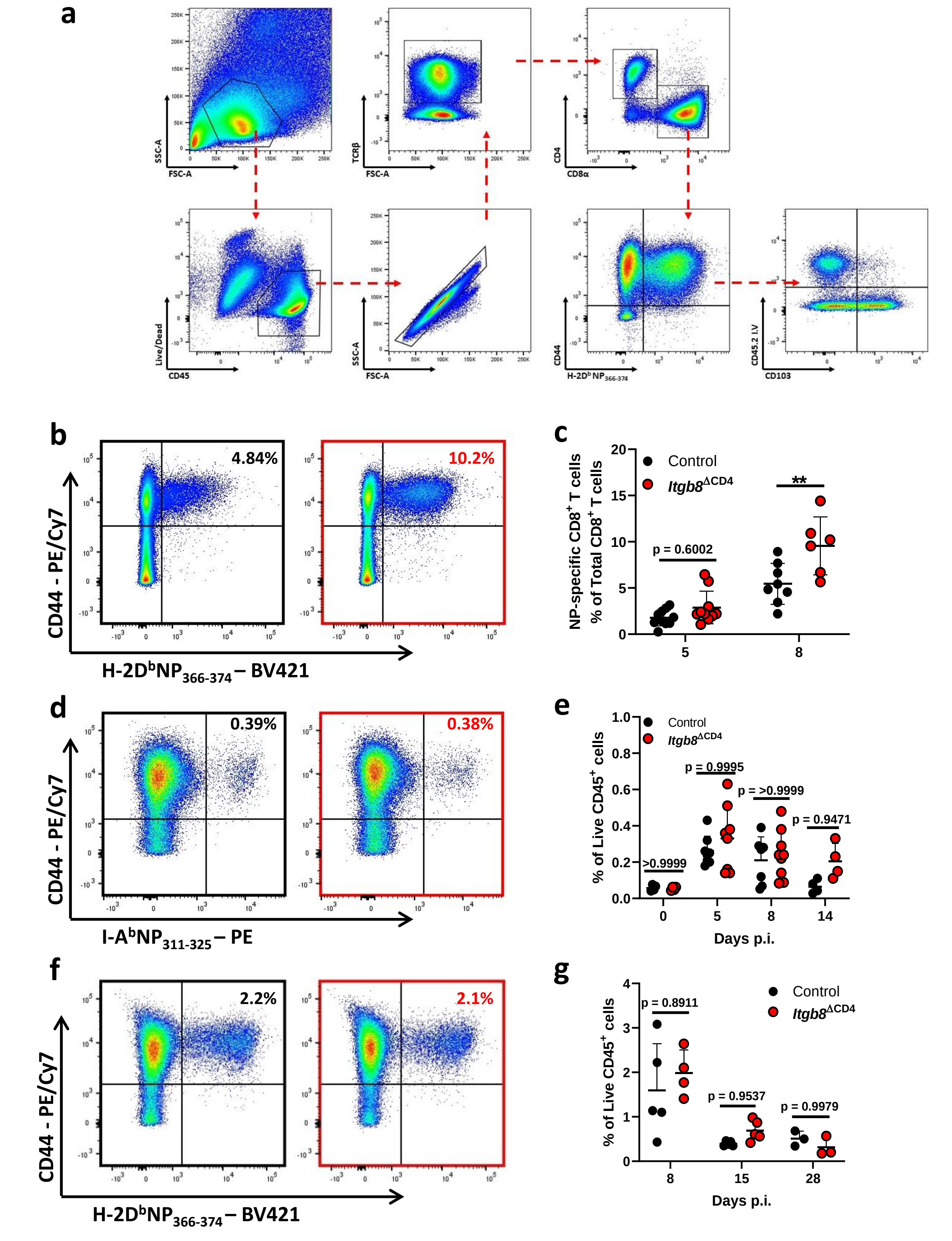


### Supplementary Figure 3

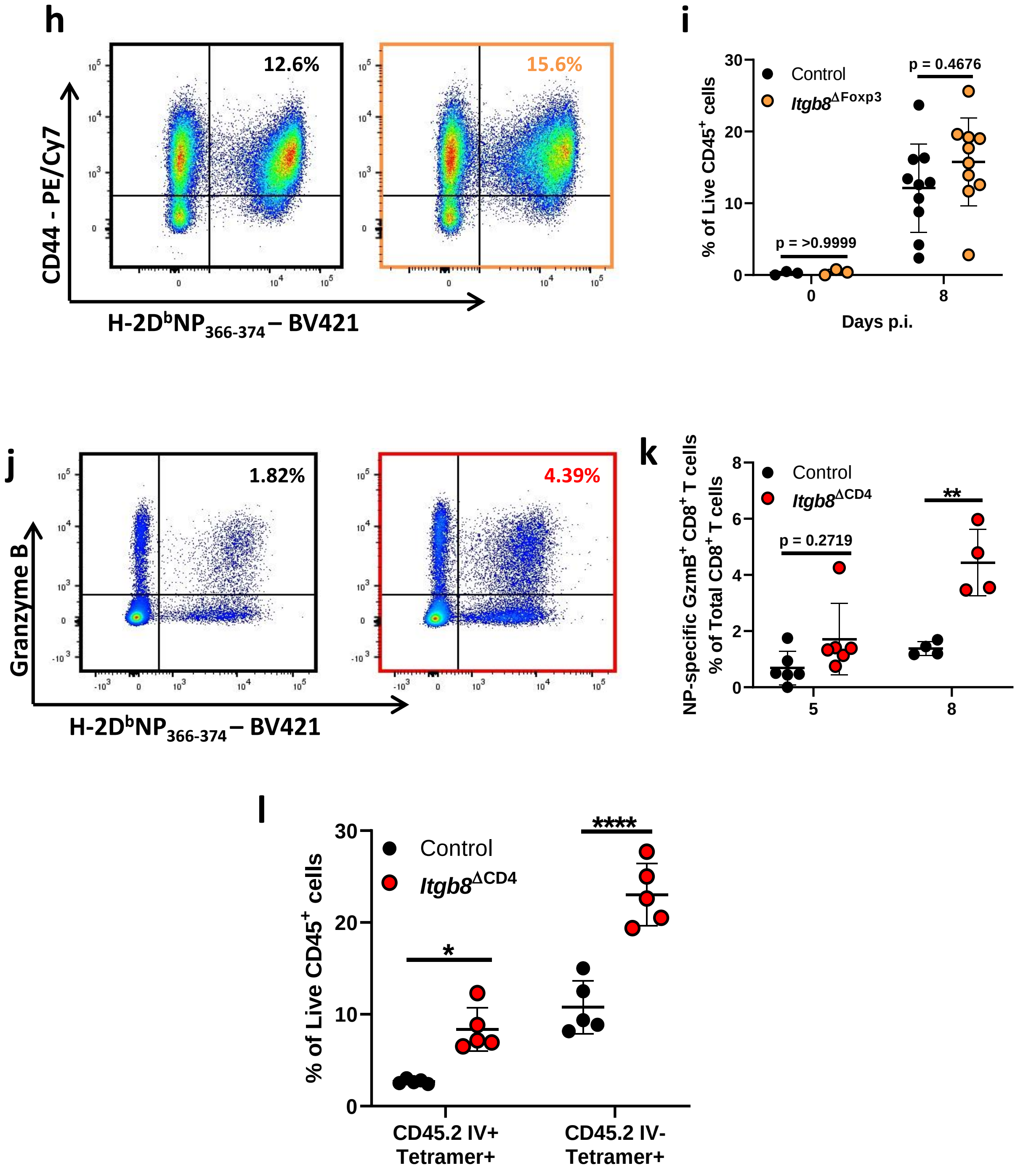


### Supplementary Figure 3 continued

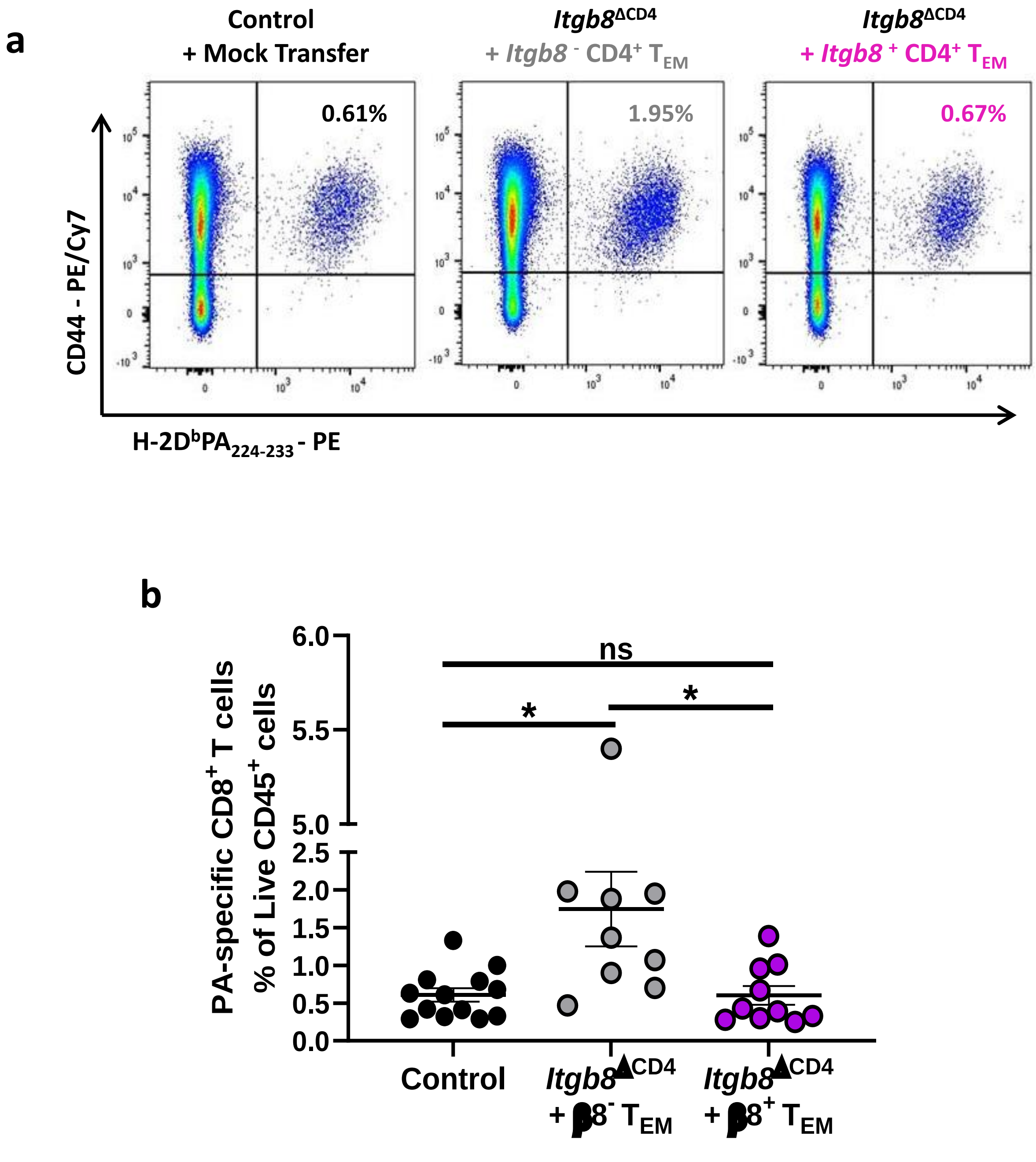


### Supplementary Figure 4

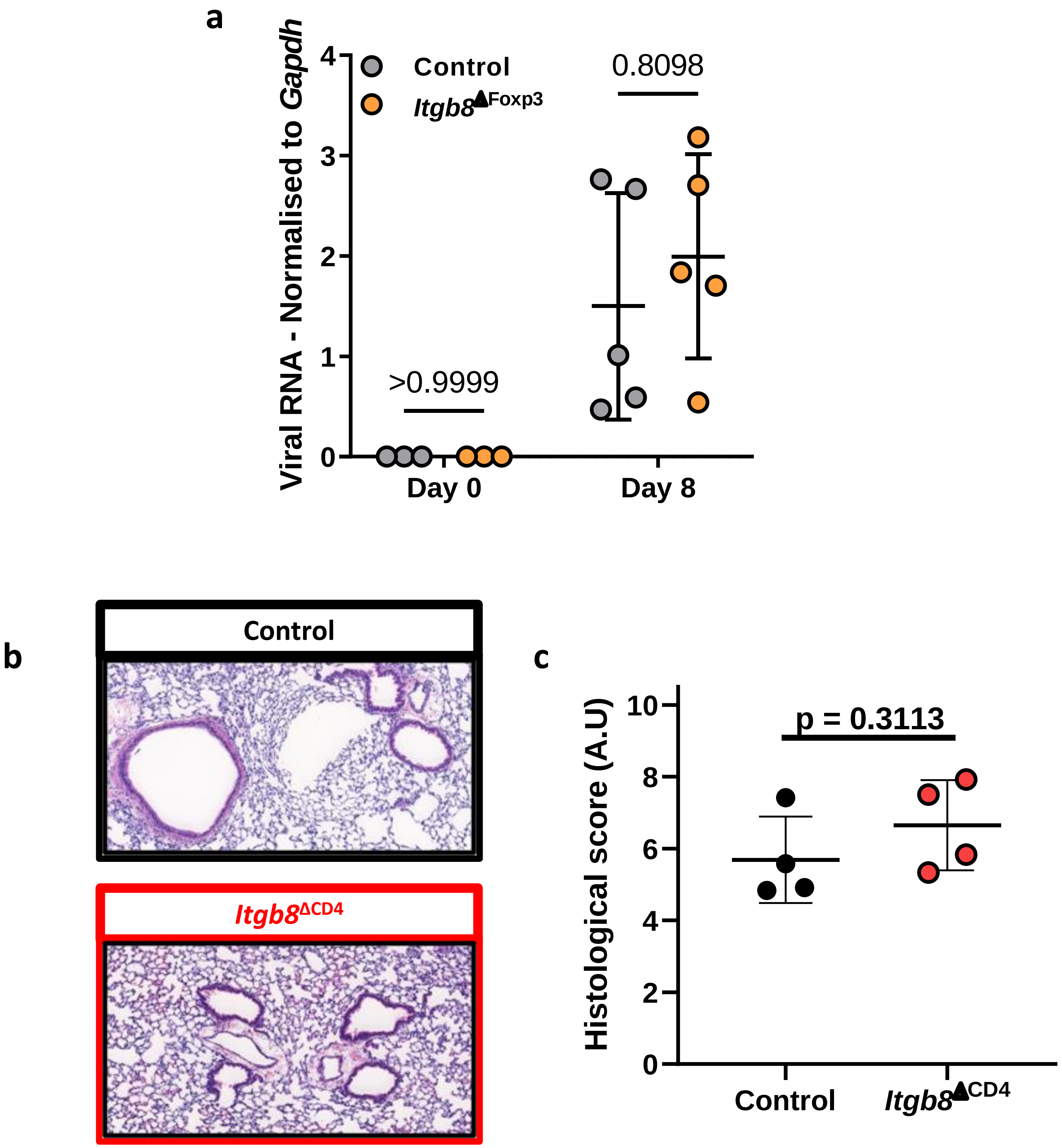


### Supplementary Figure 5

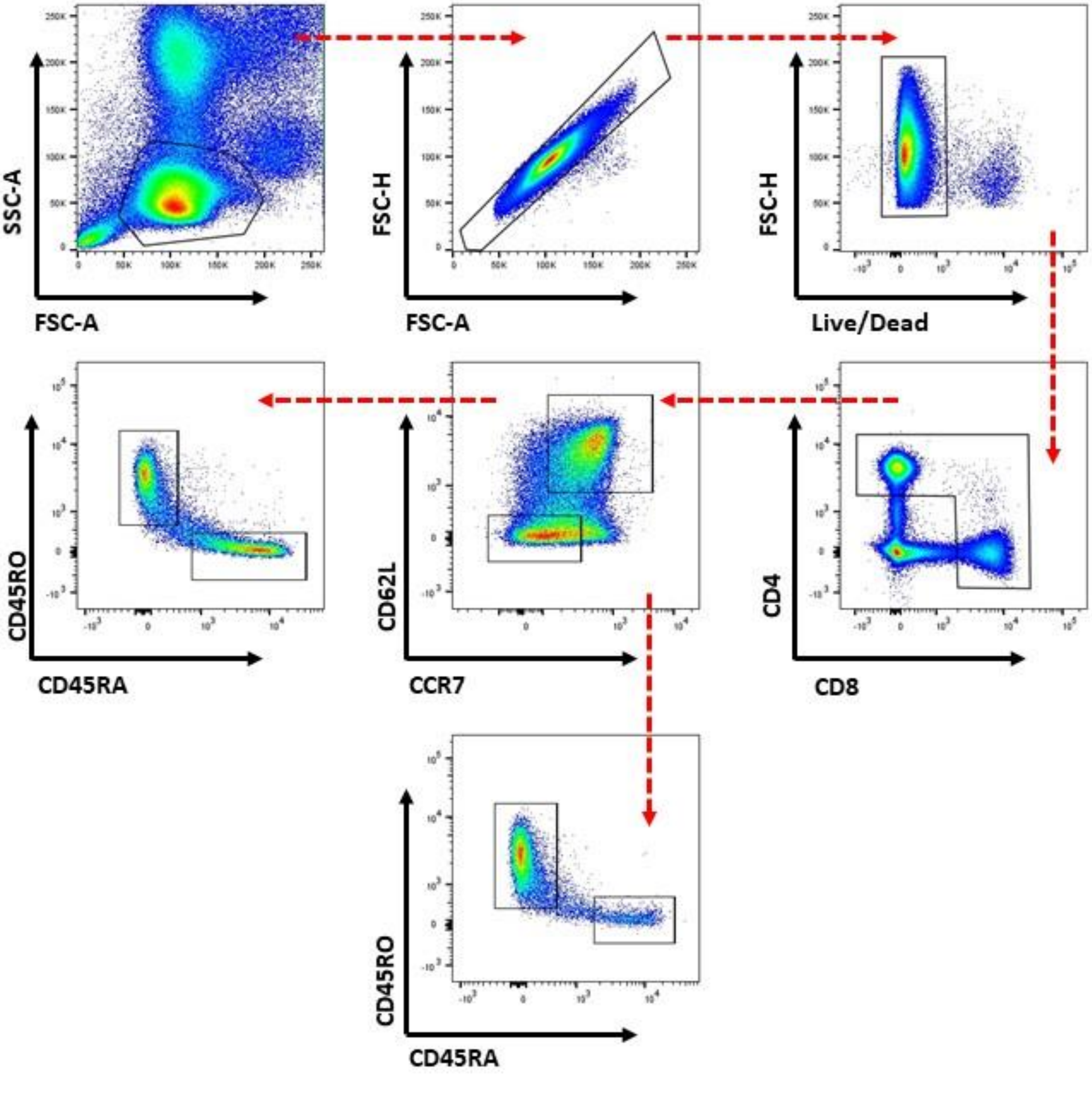


### Supplementary Figure 6
